# Are mice born normally with an egg-only diet?

**DOI:** 10.1101/2020.05.12.091660

**Authors:** Ken-ichi Isobe, Natsumi Sato, Karin Ono, Haruka Fujita, Marin kawamura, Harumi Sasaki, Saori Yamamoto, Naomi Nishio

## Abstract

Chicken eggs (egg) contain high amounts of important proteins and fat with a very low amount of carbohydrates, and also contain all vitamins and minerals needed for the chick to develop. We mated 8-week-old mice consuming an egg-only diet, and examined whether mice got pregnant and mice gave birth. Then we continued to raise newborn mice only consuming an egg-only diet compared to normal chew diet. We found that the rate of pregnancy of the mice consuming an egg-only diet was not different from that of the mice consuming normal chew diet. Further, we found that the number of pupus born from the mother consuming only eggs was not different from that of the mother consuming normal chew diet. However, the newborn mice from the mother fed only eggs did not grow healthy when they consume egg-only diet. Only few mice were alive at one month after birth. Then we change the diet just before or just after birth from egg only diet to normal chew diet and we found that both group grew healthy more than one month. These results indicate that mice are born healthy only with egg only diet.

## Introduction

Chicken egg (egg) is a conventional food that contains every nutrient needed for the normal growth of the chicken embryo. Eggs contain high amounts of important proteins and fat, with a very low amount of carbohydrates. Eggs contain all vitamins and minerals needed for chick development^1^. Chick develops independently from mother. Fetus of mammal receives nutrients from the mother via umbilical vein, whereas fetus of birds get nutrients only from albumen and yolk independent to mother.

Obesity and metabolic syndrome are currently serious problems worldwide. Type 2 diabetes, nonalcoholic fatty liver disease (NAFLD), atherosclerosis, stroke, and ischemic heart disease occur in people with those conditions^1^.

Recently, low carbohydrate diets (LCD) have been extensively investigated in several clinical states^2,3^. However, a carbohydrate-restricted diet was reported to increase mortality^4,5^.

To understand the deep biological meaning of low carbohydrate nutrition, we designed the experiments to feed mice only hen egg. Here, we asked whether murine pups could develop in the mother fed egg-only diet. Here we set up the experiments to examine the rate of pregnancy between the mice fed only eggs and the rate of healthy new born.

## Materials and Methods

### Mice

Five to eight -week-old C57BL/6 (C57BL/6n; B6) mice were purchased from SLC Japan. They were maintained on either a normal diet (ND; CLEA Rodent Diet CE-2) or on an egg-only diet, under 12-hour light and dark cycles and specific pathogen-free conditions in the Animal Research Facility at the Nagoya Women’s University, and were used according to institutional guidelines. All mice were housed up to 5 mice per cage, with ad libitum access to diet and tap water. The Animal Care and Use Committee of Nagoya Women’s University approved the study protocol.

### Mouse diets and feeding

The CLEA Rodent Diet CE-2 was the ND; it contains 8.84% water, 25.48% protein, 4.61% fat, and 61.07% carbohydrate, along with vitamins and minerals. The energy of 100g of CE-2 is 339.1 kcal (Chubu Kagaku, Japan). For the egg-only diet, we used boiled eggs. Eggs were obtained from firms in Aichi, Japan. The nutritional analysis of the boiled eggs was performed at Japan Food Research Laboratories (Nagoya, Japan). Boiled egg has a high water content (75.6g/100g), and 12.9g/100g protein, 9.6g/100g fat and 1.1g/100g carbohydrate and vitamins and minerals, which corresponds to 36.1% of calories (E) from protein, 60.8% E from lipid and 3.1% E from carbohydrates.

### Mesurement of body weight organ weight

Body weights were measured regularly and organ weights were measured at the time of sacrifice.

### Biochemical analysis

Serum samples were obtained from the mice at the time of sacrifice. Serum glucose, ketone bodies were measured by enzymatic colorimetric assays. Blood glucose was measured by using the LaboAssayTM Glucose kit (Wako Pure Chemicals, Osaka, Japan). Ketone bodies, including total ketone bodies (T-KB) and 3-hydroxybutyrate (3-HB) were measured by AUTO WAKO Total Ketone Bodies and 3-HB (Wako Pure Chemicals, Osaka, Japan). Wavelengths were measured with an xMark™ Microplate Absorbance Spectrophotometer (Bio Rad).

### Statistical analyses

Data are expressed as means□±□standard deviations (SD).

## Results

### 1. The number and body weight of the new born mice from the mother fed only eggs were almost same as those of the mother fed chew diet

In order to examine whether the mice fed only eggs give birth, we set up the experiments of mating the mice in a cage with one male and five or six female mice consuming only eggs or consuming control diet (CE-2). Among 11 female mice 9 mice became pregnant in egg-only diet in one month (82%). Among 10 mice 8 mice became pregnant in control diet (80%). These indicate that the rate of pregnancy was not different between egg-only diet and control diet. Further, average number of newborn mice from the mother fed only eggs was 6.2, whereas average number of newborn mice from the mother fed control diet was 7.0 (Fig. 1a). We found that the appearance of the newborn mice from the mother fed only eggs was healthy. The average body weights of newborn mice from the mother fed only eggs was 1.29g, whereas the average body weights of newborn mice from the mother fed control diet was 1.50g (Fig.1b). This means that newborn mice born from the mother fed egg-only diet were slightly lower than the mice born from the mice fed control diet.

**Figure 1.**
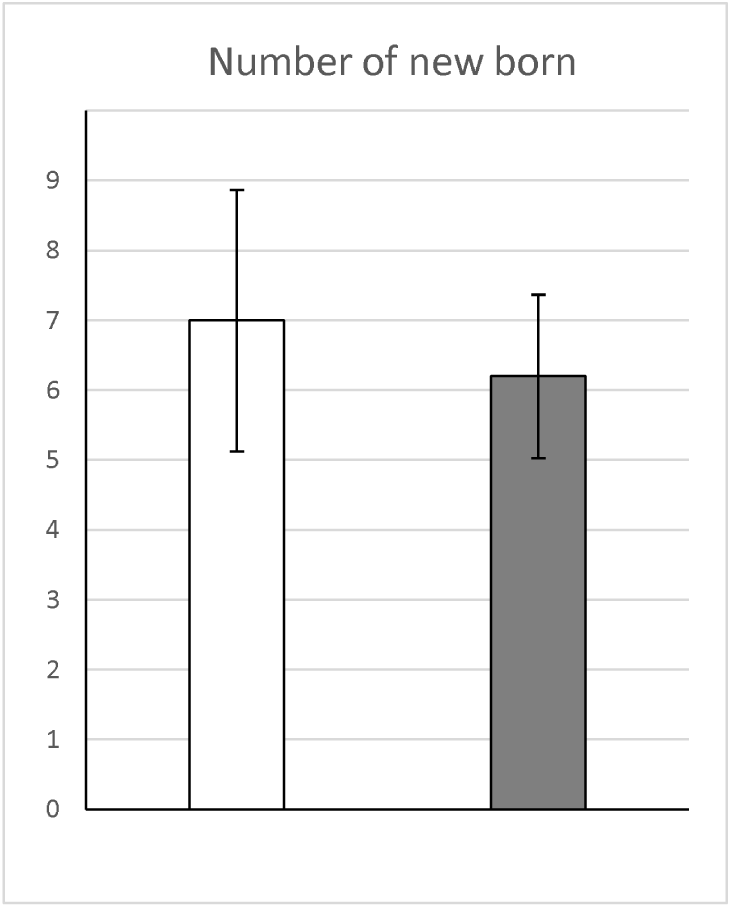
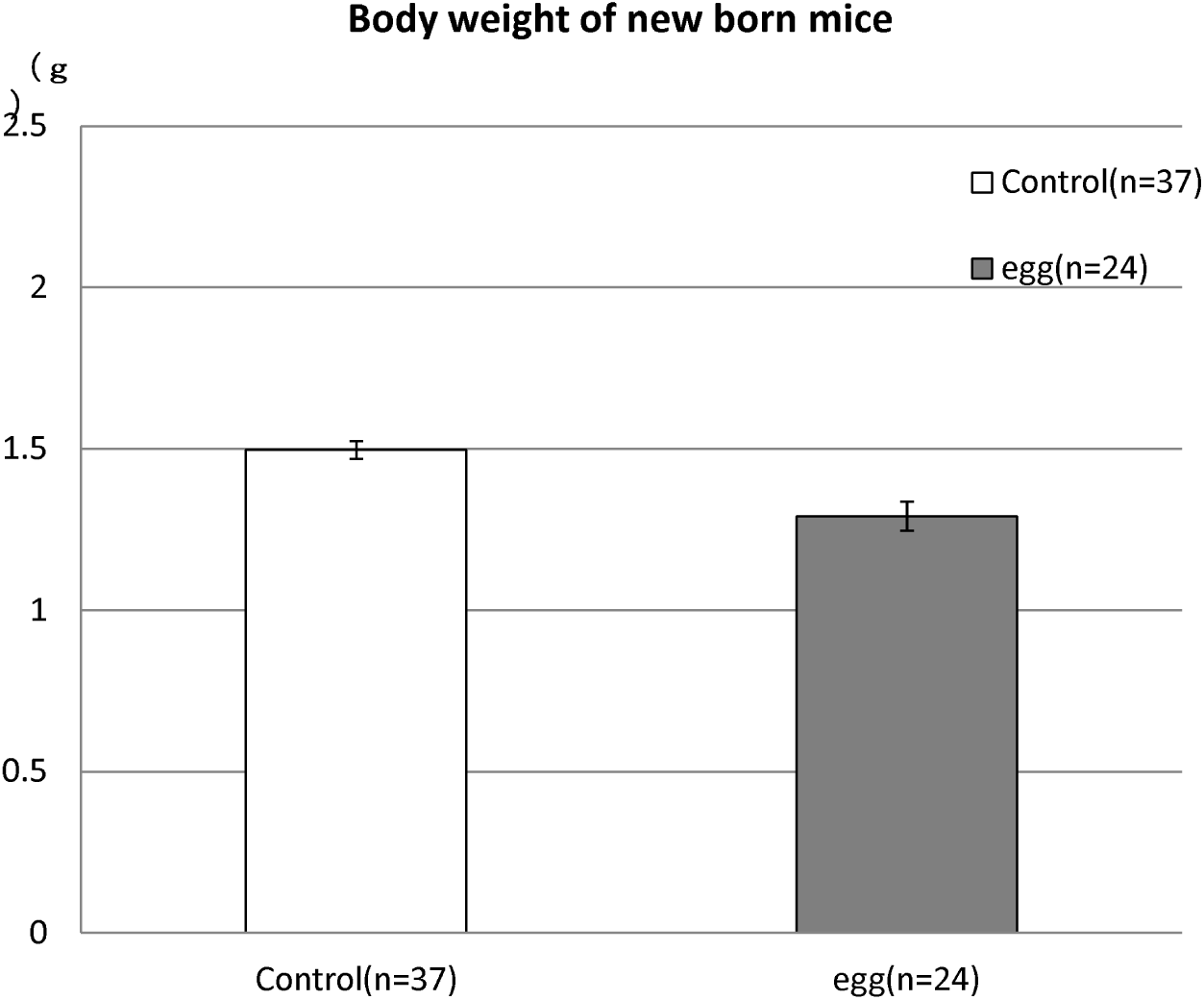
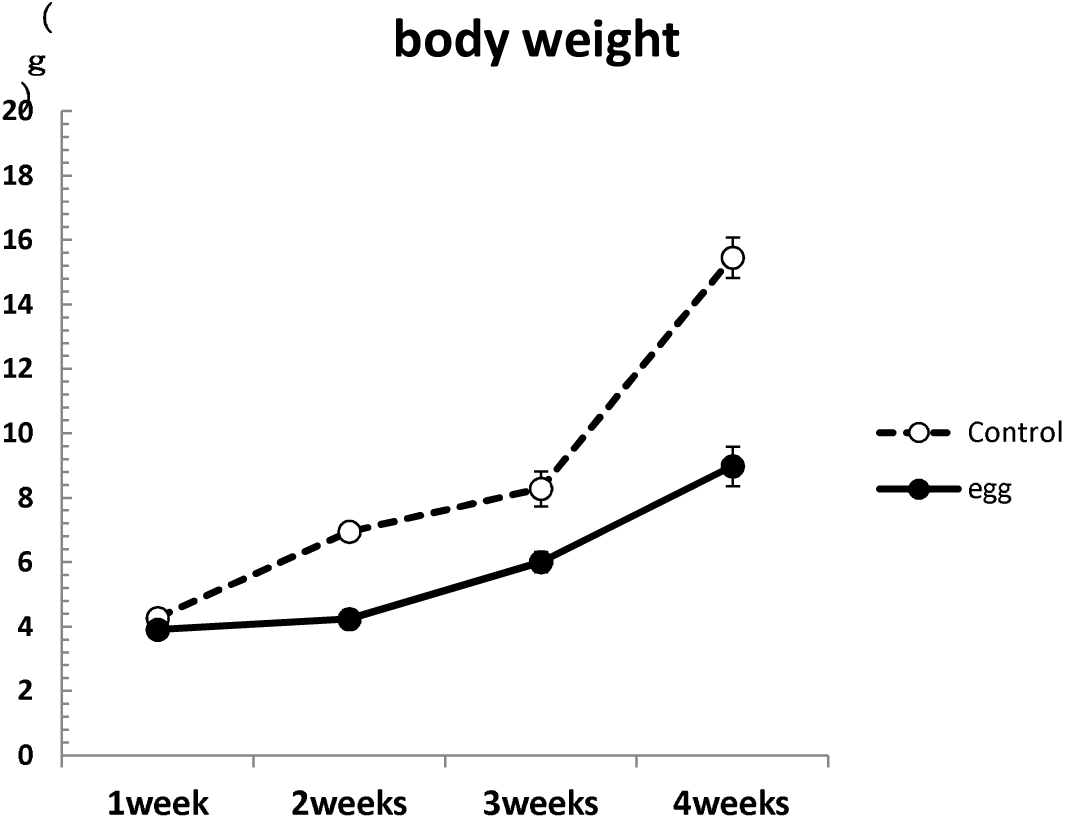
The number and body weight of the newborn mice from the mother fed only eggs. Eight-week-old one male mice and five or six female mice were housed together in a cage. They were fed 1) only CE-2 (control) or 2) only eggs, ad libitum. Pregnant mouse was moved to a new cage and fed the same diet. (a) Number of mice born from the mother fed normal diet or egg-only diets. (b) Body weights of the newborn mice from the mother fed CE-2 (control) or egg-only diet were measured at birth. Data shown are the mean ratios ± standard deviations (control n=37, egg n=24). (b) Body weights of newborn mice nursed by the mother fed CE-2 (control) or egg were measured at 1, 2, 3, 4 weeks.

### 2. The newborn mice from the mother fed only eggs did not grow healthy when they consume egg-only diet

We continue to breed newborn mice by same diet. At one week they were healthy and body weights were almost the same as the newborn mice from the mother fed control diet. However, at two weeks they were smaller than those born from the mother of normal diet. The weights of liver of 2, 3, 4 weeks of the mice born from the mother egg-only diet were lower than those born from the mother of normal diet (Fig.1c).

### 3. The mice bred by ND just before or after birth from egg-only diet grow healthy

In order to know whether the mice born from egg-only mother can not survive by milk feeding, we changed the diet just before or just after birth from egg only diet to control diet (Fig.2a). Almost all mice of second group (egg to control diet before birth) and the mice of third group (egg to control diet after birth) survived more than four weeks, whereas the mice of third group died after 1 week and many mice were dead at 4 weeks (Fig.2b). At birth the number of mice of all groups were almost same (Fig.2c) and looked healthy (Fig.2d). At birth body weight of all groups were not different (Fig.2e). Body weights of second and third group (egg to control diet before birth and after birth) were not different from fourth group (control diet) until four weeks, although the body weights of first group (egg-only diet) were much lower than those of other groups (Fig.2e). By autopsy we measured liver and heart weight from 1 week to 4 weeks. We found that liver weight of all groups were almost same at 1 week. Liver weights were much lower in only first group (egg-only mother) from 2 weeks to 4 weeks than those of other groups (Fig.3a). On the contrary heart weights of the mice bred by the mother of first group (egg-only diet) were higher than those of other group (Fig.3b). Especially heart weight/ body ratio of the mice bred by mother of egg-only diet was much higher than those of other group (Fig.3c). This means that changing the diet from egg to control diet at birth rescue the mice from some problems of the milk of the mother bred egg-only diet.

**Figure 2.**
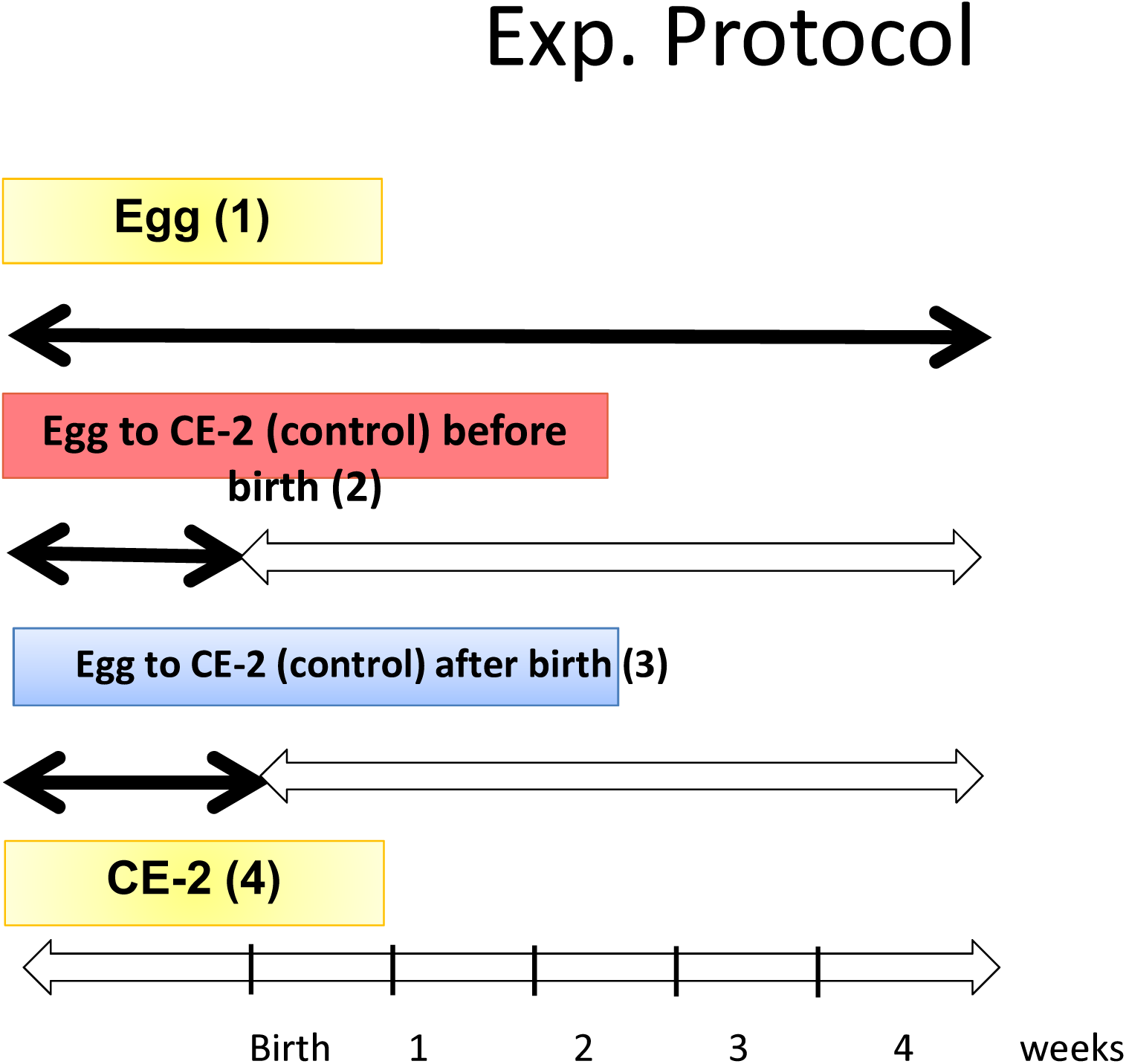
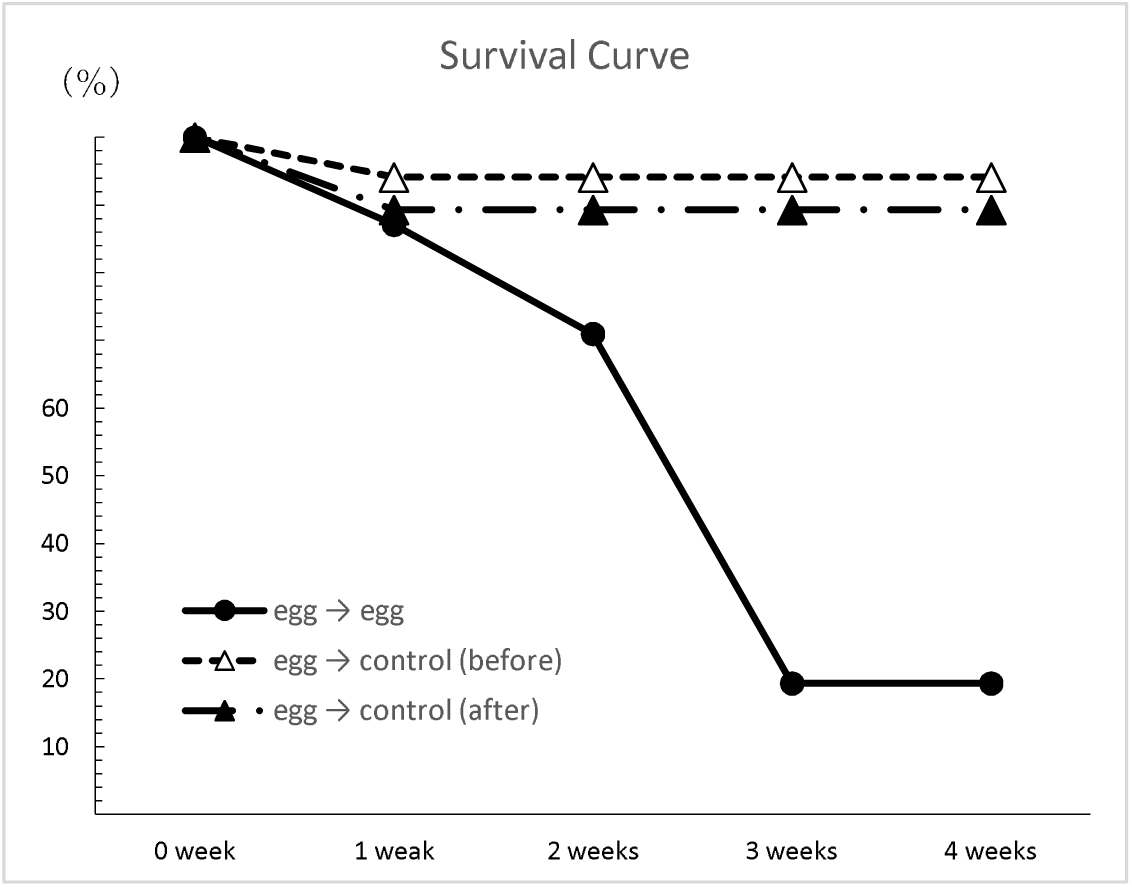
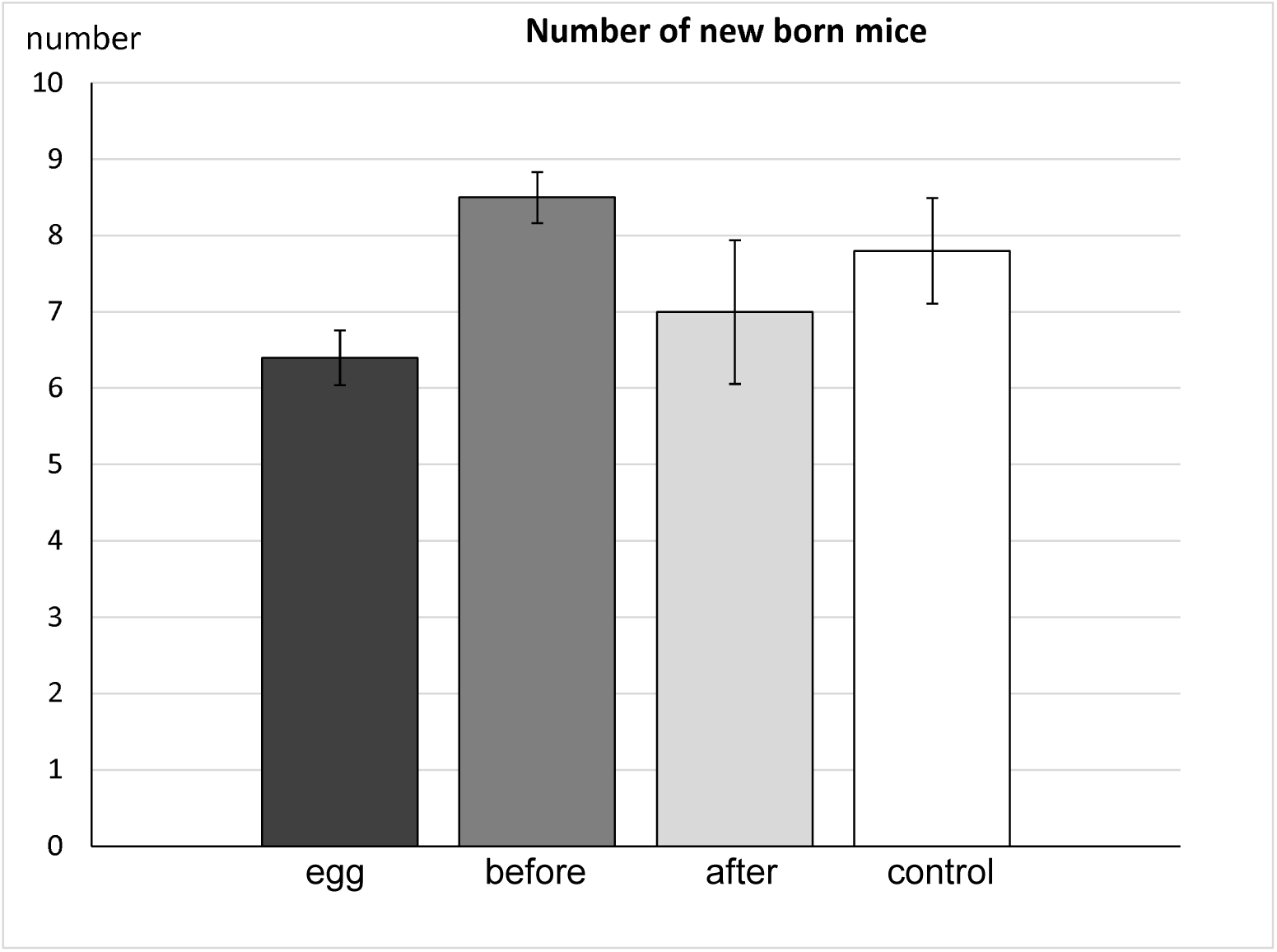
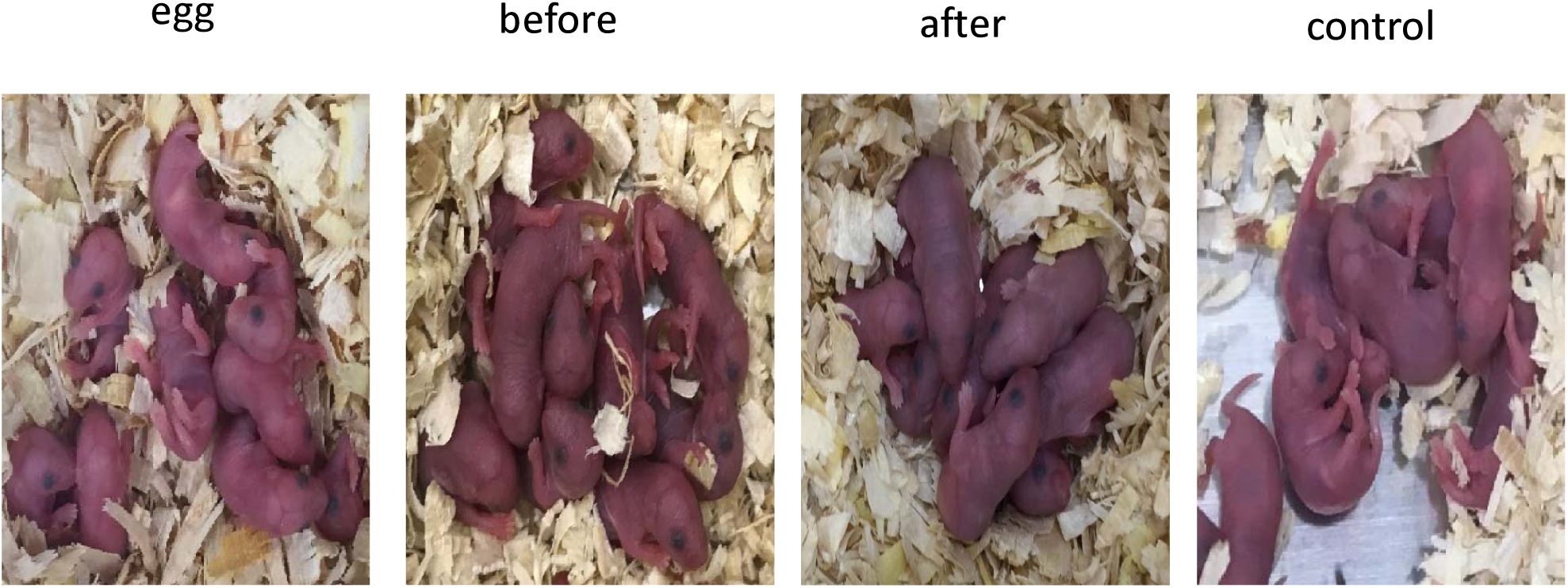
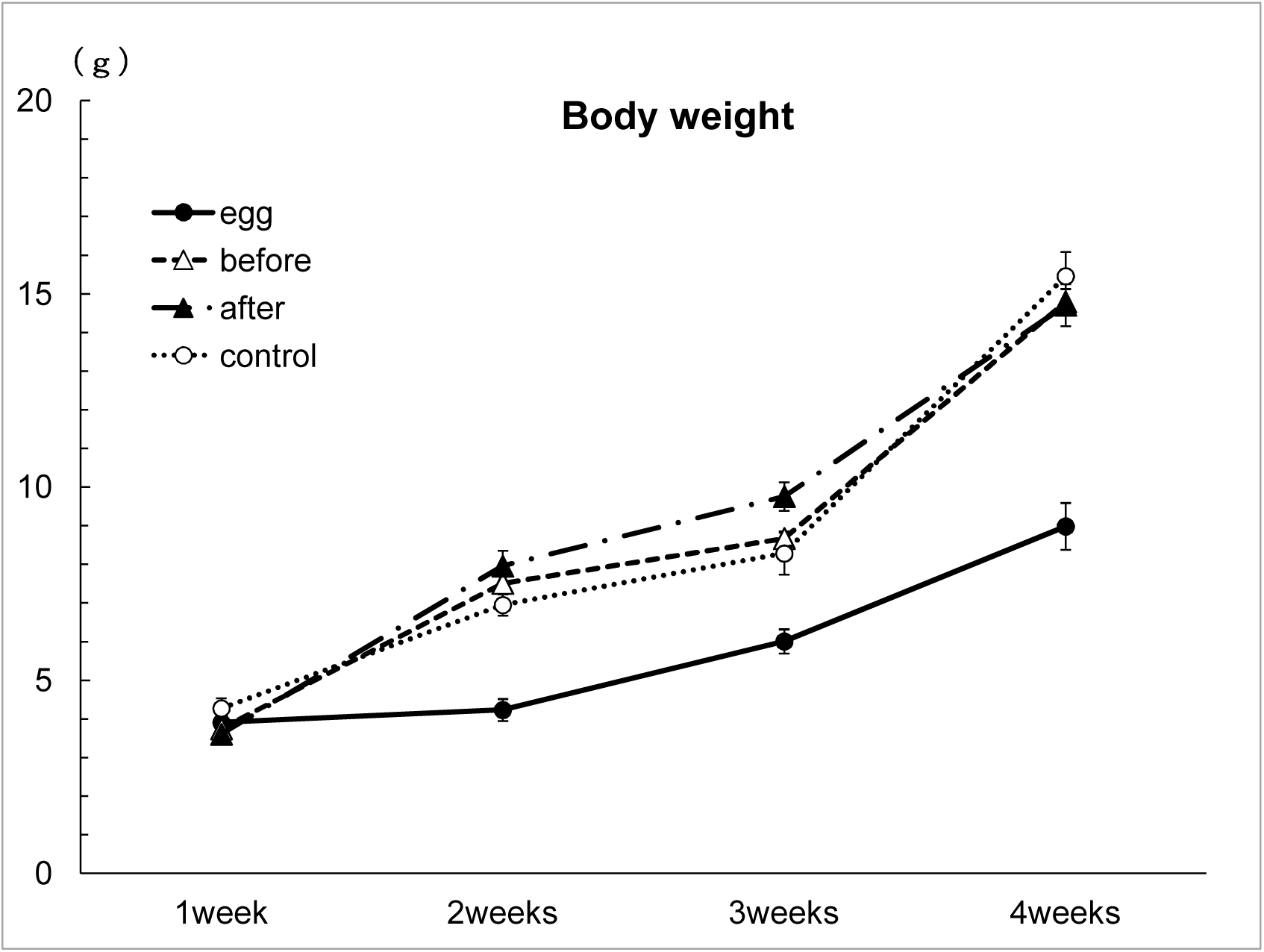
The mice bred by control diet just before or after delivary from egg-only diet. Pregnant mice fed only eggs were bred by 1) egg-only diet, 2) CE-2 (control) diet just before delivery or 3) control diet just after delivery. 4) Pregnant mice fed control diet were continued to be bred by control diet after delivery. (a) Experimental Protocol. (b) Survival curve. Number of mice alive at 0,1,2,3 and 4 weeks. (c) Average number of newborn mice from one female of each group. (d) Photographs of newborn mice of each group. (e) Average body weights of mice of each group at 1, 2, 3, and 4 weeks. Data are expressed as means□±□standard deviations.

**Figure 3.**
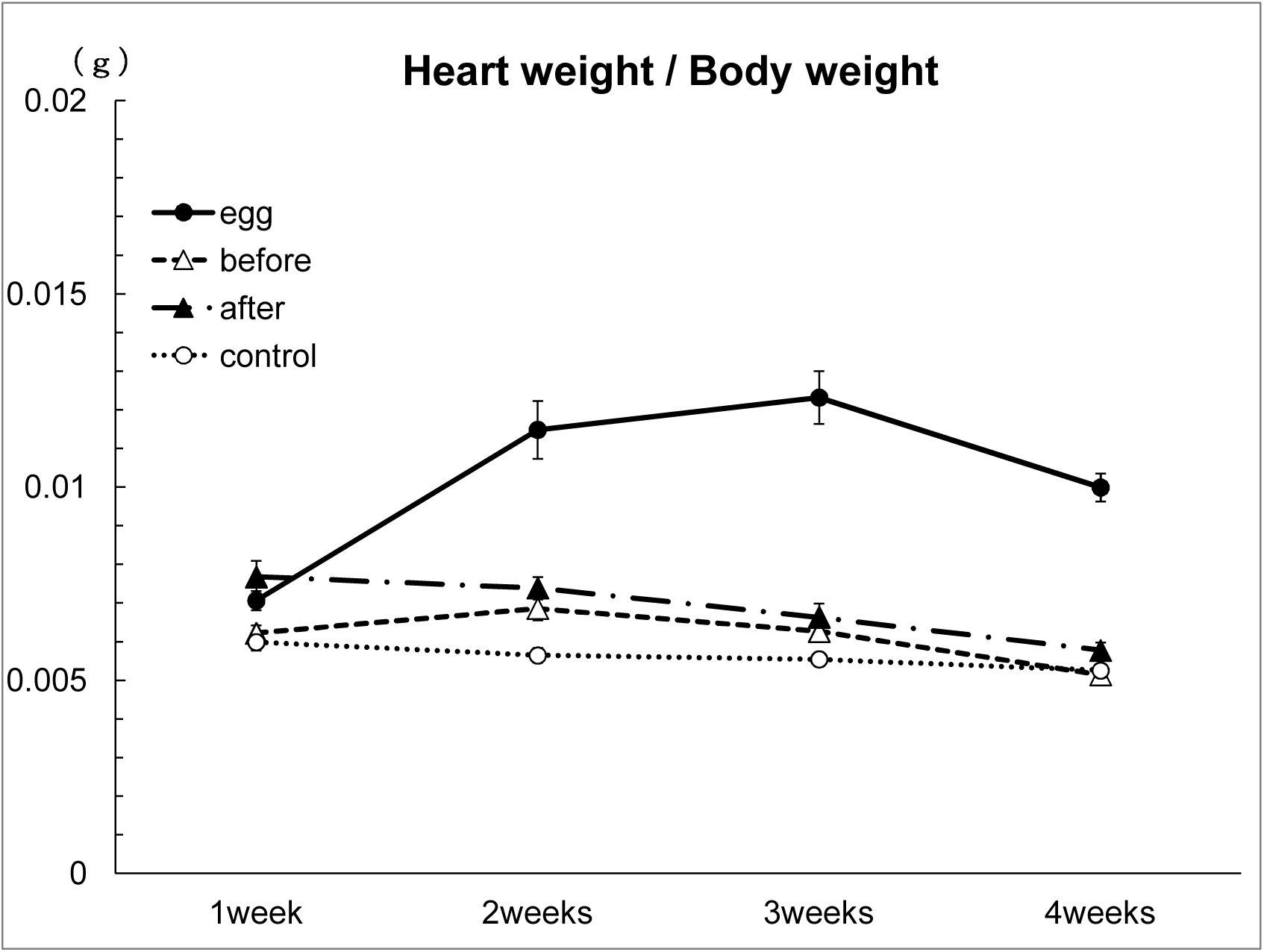
Organ weights of the mice bred by control diet just before or after delivary from egg-only diet. (a) Liver tissues of each group were removed from the mice used in Figure 2 and liver weights were measured. (b) Heart tissues of each group were removed from the mice used in Figure 2 and heart weights were measured. (c) Heart tissue weights were divided with 100x body weights of the same group. Data are expressed as means□±□standard deviations.

### 4. Keep the glucose level and high ketone bodies in egg-only pregnant and nursing mice

In order to know the gluconeogenesis of the pregnant and nursing mice fed only eggs, we measured the blood glucose around 13^th^ to 16^th^ of pregnant mother. We found that pregnant mice fed only eggs had almost the same level of blood glucose as those of pregnant mice fed control diet (Fig.4a). In nursing mice, the blood glucose levels of the mother fed only eggs were almost the same as those of the mice fed control diet. However, the blood glucose levels of the mother fed control changed from eggs after birth were slightly higher than those of the mice fed only eggs (Fig.4b).

**Figure 4.**
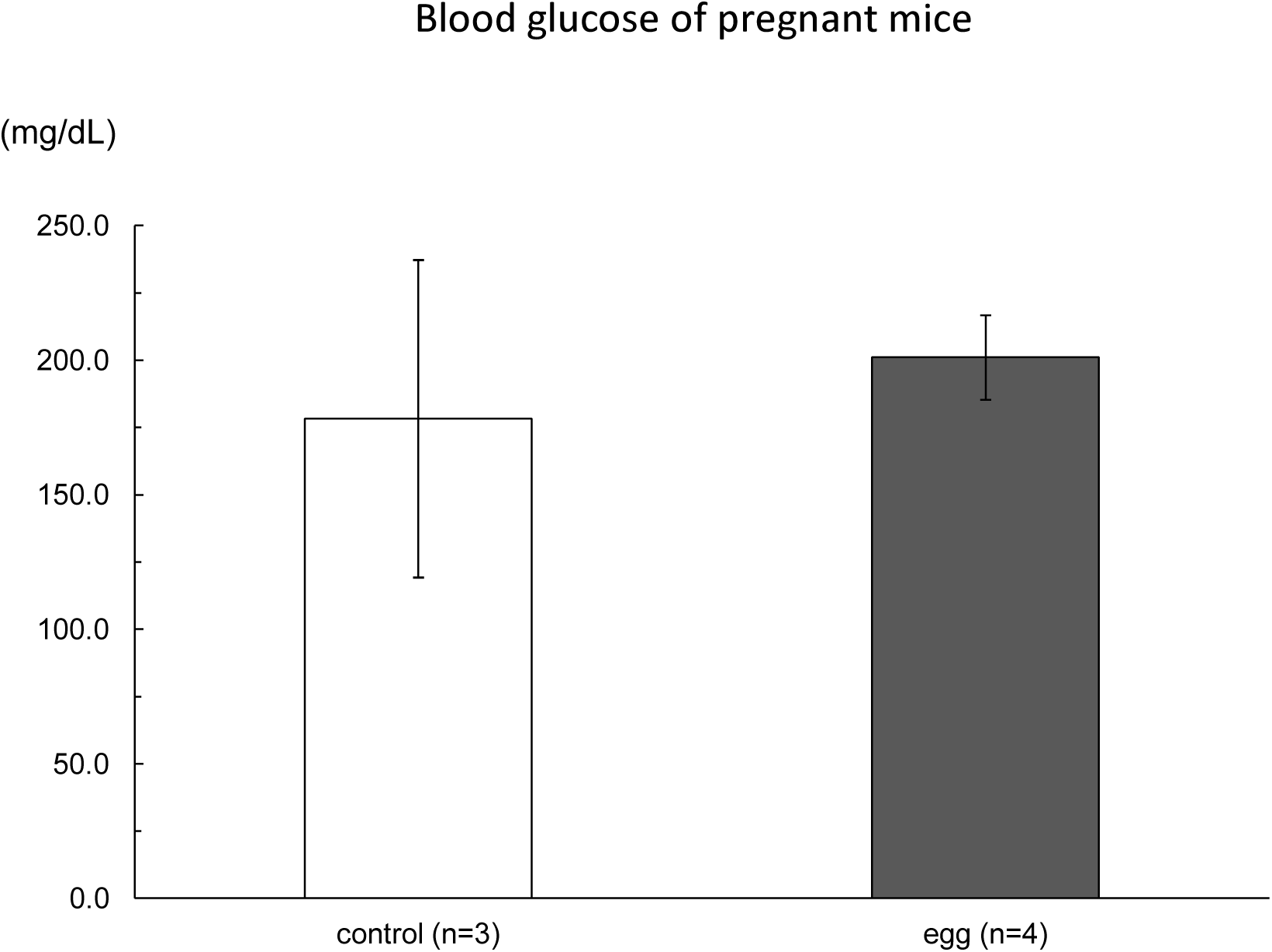
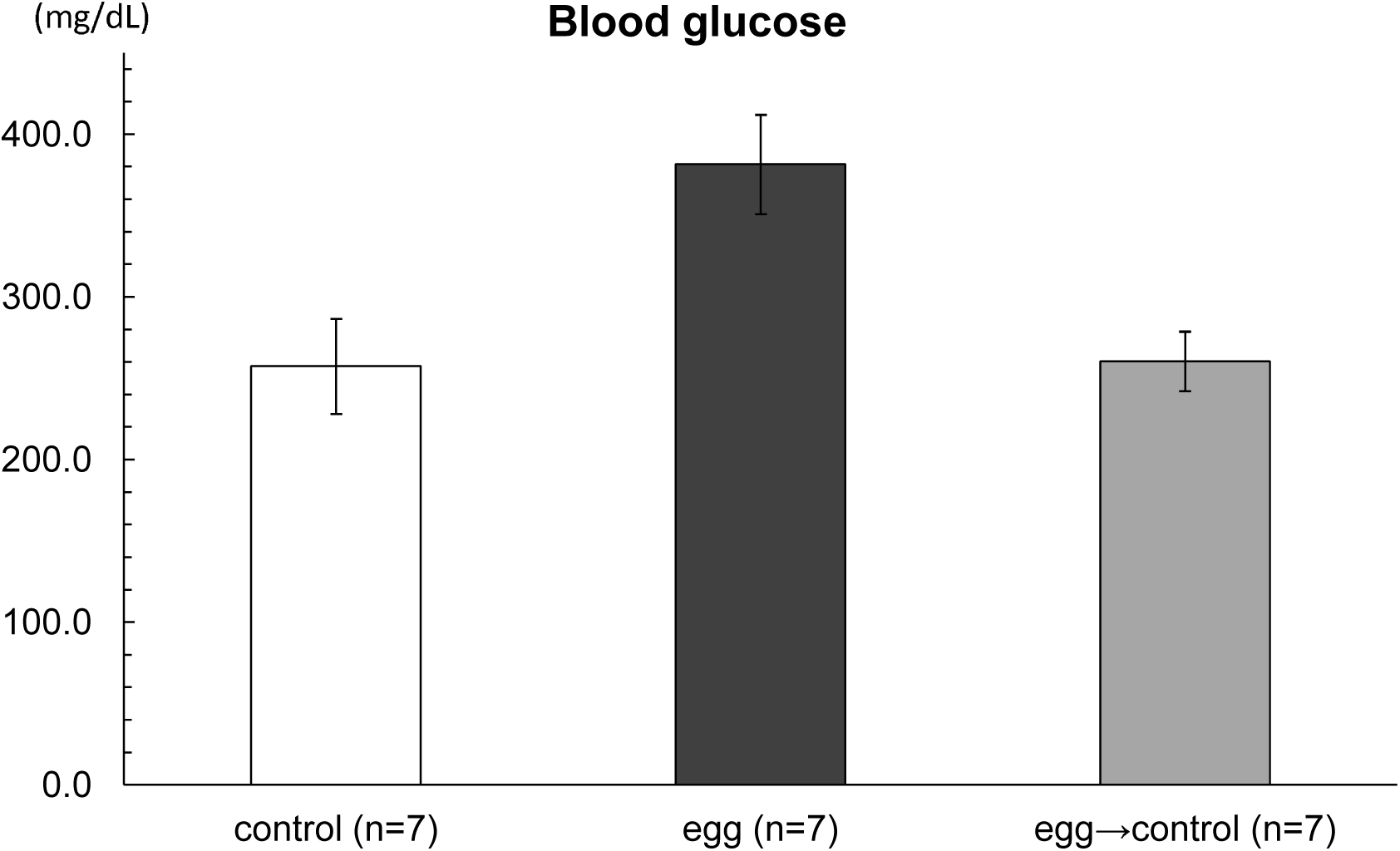
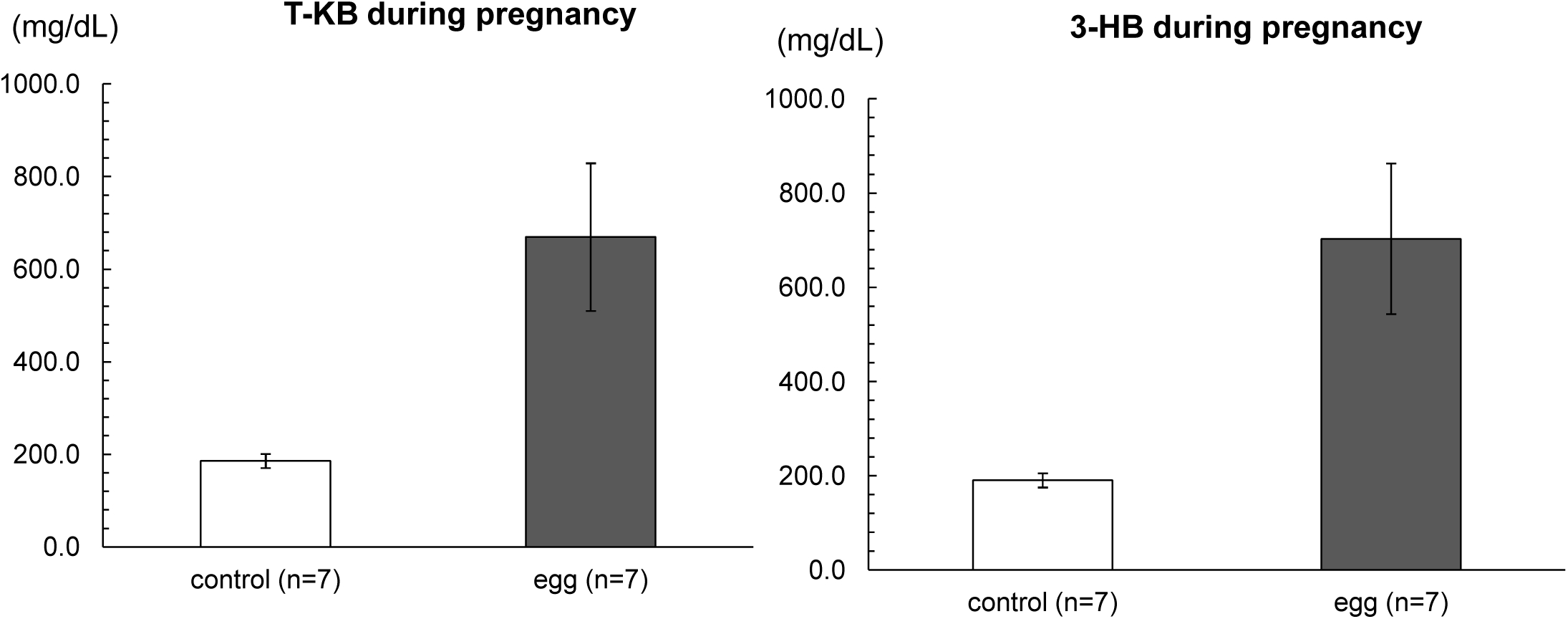
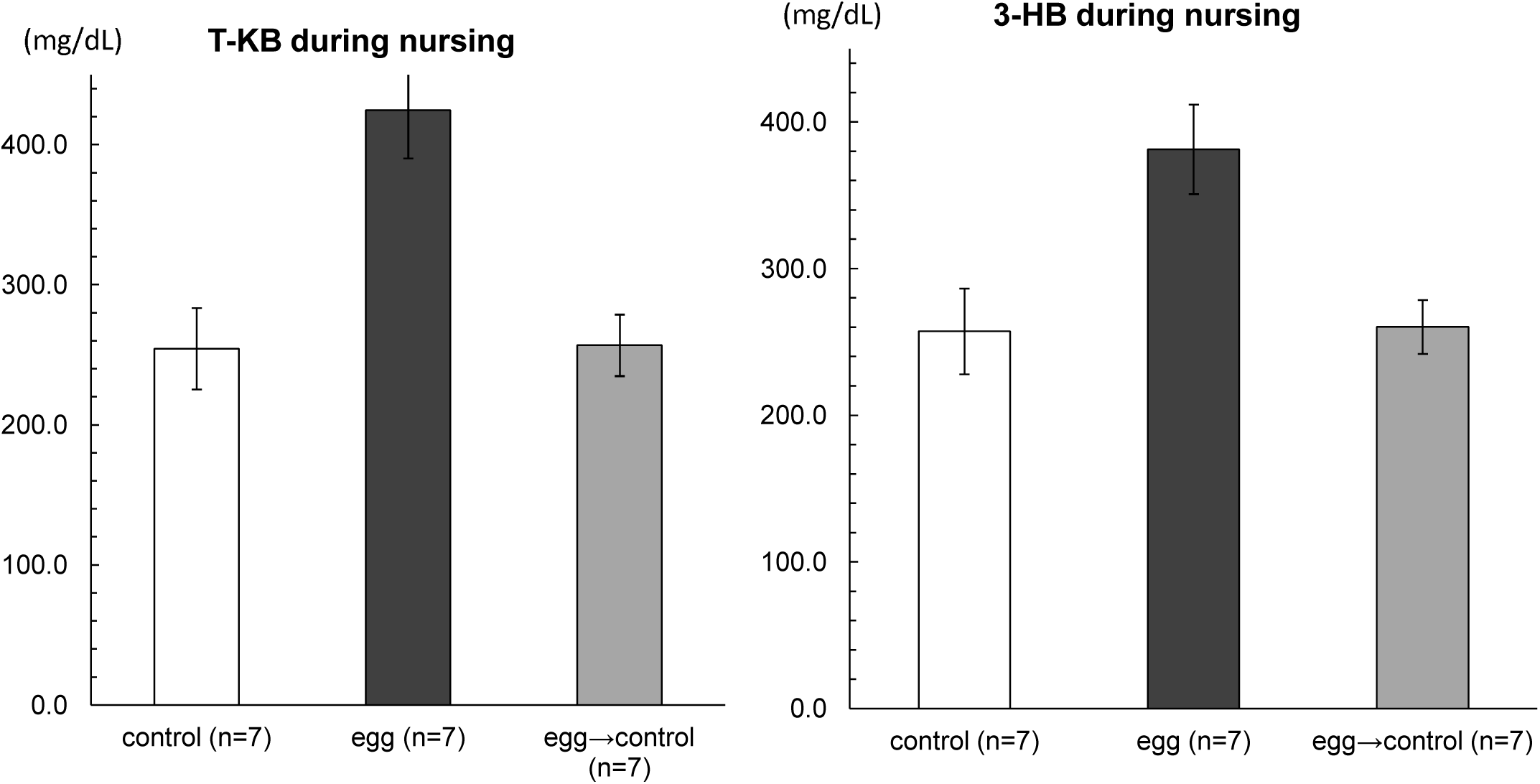
Keep the glucose level and high ketone bodies in egg-only pregnant and nursing mice. (a,c) Serum from the around 13^th^ to 16^th^ of pregnant mice bred only eggs or normal diet were taken in the morning and levels of the serum glucose (a) or levels of the serum T-KB and 3-HB were measured. (b,d) Serum from each group of the mice in Fig.2 were taken in the morning, and levels of the serum glucose (c) or levels of the serum T-KB and 3-HB (d) were measured. Data are expressed as means□±□standard deviations.

The ketone bodies in the serum of the egg-only pregnant and nursing mice were examined and compared with those of the control diet mice. Both T-KB and 3-HB were much higher in the serum of the egg-only pregnant mice than in the serum of the control pregnant mice (Fig 4c). Both T-KB and 3-HB were higher in the serum of the egg-only nursing mother mice than either serum of normal diets of nursing mother mice or serum of egg to normal diet of nursing mother mice (Fig.4d).

## Discussion

### 1. The mice were born healthy from the mother fed only eggs

In this paper we have shown that female mice fed only eggs can successfully deliver term babies. The new born mice are healthy, because almost all new born mice from egg-only mother are healthy and gain same weight at 1 months after birth as those of the mice born from the mother fed ND when they were nursed with the mother fed ND. Because, chick and mammal use liver as a major organ of gluconeogenesis, they need glucose at the first stage of development until functional liver is developed. Glucose is used as an energy and a source of nucleic acids (through pentose phosphate pathway) for quickly dividing tissue cells^6^. The albumen was made up of 82% water, 17% protein, and 0.5% carbohydrates, whereas the yolk was made up of 53% water, 15.6% protein, 27.4% fat, and less than 0.5% carbohydrates^7^. In the studies of nutritional need of chick development, it has been demonstrated that carbohydrates are used for the initial phase of chick development as a source of nucleic acids for quickly dividing cells. Then at middle phase of chick development proteins are mainly used. The last phase of chick development (after 14^th^ days of incubation) fats are used for energy source as □-oxidation^8,9^. Chicken embryos grow exponentially during the final week of incubation, with about 90% of the total energy for growth derived from β-oxidation of yolk fatty acids^10^.

Embryonic development of mice is different from that of chicken. Mice get energy from umbilical vein, which contain high amount of glucose for energy. Human fetuses receive nutrients from the mother, but after birth, humans receive nutrients in different ways depending on their regional habits or religion.

### 2. High keton bodies in egg-only pregnant

We have shown that the levels of T-KB and 3-HB were more than 3 times higher in the egg-only pregnant mice than in normal diet mice. In nursing those levels were also higher in the egg-only nursing mice than in normal diet mice. In mammals, ketone bodies are produced predominantly in the liver from FAO-derived acetyl-coenzyme A (CoA), and they are transported to extrahepatic tissues for terminal oxidation. This physiology provides an alternative fuel that is augmented by relatively brief periods of fasting, which increases fatty acid availability and diminishes carbohydrate availability^11^. Multiple studies suggest diverse signaling roles for ketones, even in carbohydrate replete states, a subset of which could be independent of their metabolism ^12^.

### 3. Controversial clinical studies of low-carbohydrate and high fat diet in gestational diabetes Mellitus

Gestational diabetes Mellitus (GDM) is one of the urgent medical problems in the world. The management of GDM still poses many unanswered questions^13^. The restriction of dietary carbohydrate (CHO) has been started for the treatment of GDM at a century ago ^14^. Meta-analysis of low-CHO for treatment of GDM has been done recently^15,16^. These analysis indicated that many studies of low-CHO for treatment of GDM indicated a larger decrease in fasting and postprandial glucose associated with lower infant birth weight. However, they critically evaluated the many low-CHO studies as low to very low quality^16^. Further low-CHO with high lipids increased fasting total cholesterol and FFA, which increased inflammation and insulin resistance^17^. WHO guideline recommended a low-fat diet (<30% of energy) and limiting saturated fatty acids to less than 10% of energy intake by replacing them with unsaturated fatty acids^18^. WHO guidline was based on the studies of North America and Europe, where people consume an excess energy with a low carbohydrate^19^. However, outside these regions especially in Asia most people consume very high carbohydrate diets such as refined rice. Recently several epimediological studies showed controversial results. Increased dietary fat intake reduced mortality and was not linked to cardiovascular disease ^20^. Increased dietary fat intake reduced type 2 diabetes and cancer (21). Intake of dietary saturated fatty acids (SFAs), which has long been perceived as unhealthy, was not associated with increased risk of insulin resistance or cardiovascular disease ^22,23^. More recent study showed that people having diets comprised 64E% fat, 16E% protein, and 20E% carbohydrate for 6 weeks with stable body weight, insulin/glucosehomeostasis in the fasting state was changed in a non-diabetic direction, as fasting plasma insulin concentration decreased after high-fat diets. Same results were obtained by saturated and unsaturated fats^24^.

### 4. A model of the study of low carbohydrate diet for the treatment of gestational diabetes

There exist several studies of the animal model of GDM. Exposure to high–saturated-fat diet (HFD) before 1 month and during gestation has been shown to induce GDM. Glucose metabolism and HFD-associated placentral oxidative stress have been shown to the cause of GDM^25,26^. In our model of egg-only diets C57BL/6 mice are not obese during pregnancy, whereas mice become obese during pregnancy in HFD.

We presented the results here that pupus born from the mother consuming only eggs looked healthy and they grow normally when they were bred by normal diet in nursing. Our results indicate that hen eggs having low-CHO and high fat are safe for murine mother.

## Acknowledgments

This research was supported by 16H05280 and 19K11687 of the Ministry of Education, Science, Technology, Sports and Culture, Japan.

